# A High-Sensitivity and High-Throughput LC-MS Workflow for Low-Input and Single-Cell Proteomics

**DOI:** 10.64898/2026.10.26.762077

**Authors:** Yusei Okuda, Shio Watanabe, Yusuke Endo, Ryo Konno, Hiromasa Mitsui, Daisuke Nakajima, Yoshio Kodera, Osamu Ohara, Daisuke Higo, Yusuke Kawashima

**Author notes:** Corresponding Author Yusuke Kawashima.

## Abstract

Single-cell proteomics (SCP) requires sample preparation that minimizes sample loss and highly sensitive LC–MS analysis because the amount of protein available from a single cell is extremely limited. Here, we developed an integrated workflow for low-input and single-cell proteomics based on single-cell proteomics by LMNG-assisted sample preparation (scpLASP), which enables microliter-scale sample preparation using standard laboratory consumables. Systematic optimization of peptide reconstitution conditions identified 0.01% n-dodecyl-β-D-maltoside (DDM) combined with 0.01% lauryl maltose neopentyl glycol (LMNG) as the condition yielding the highest number of identified peptides among those tested for low-input samples. Optimization of the LC mobile-phase composition identified 0.5% acetic acid with 1% dimethyl sulfoxide (DMSO) as the optimal condition, increasing the mean number of identified protein groups from 6,767 with 0.1% formic acid to 7,409 for 250 pg HeLa digests analyzed using an Orbitrap Astral Zoom. Shortening the LC gradient from 11 to 5.5 min enabled higher analytical throughput while maintaining mean identification counts of 6,558 and 7,088 protein groups from 100 and 250 pg HeLa digests, respectively, with median protein-level coefficients of variation of 9.20% and 6.99%. When the optimized workflow was applied to HEK293-derived Expi293F single cells using the same 5.5-min LC gradient, an effective analytical throughput of 135 samples per day was achieved with an injection volume of 4.5 µL, accounting for sample-loading time. This workflow identified a median of 5,655 protein groups per cell using an empirical spectral library generated from 20-cell samples. Overall, this study establishes a sensitive and high-throughput workflow for low-input and single-cell proteomics that combines low-loss microliter-scale sample preparation with optimized LC–MS analysis, achieving high proteome depth and analytical throughput without requiring specialized nanoliter-scale sample-preparation devices.

## 1. Introduction

Single-cell proteomics (SCP) has rapidly emerged as an important technology for characterizing cellular heterogeneity at the protein level (1, 2). Unlike transcriptomic analysis, proteins cannot be amplified in the same manner as nucleic acids. Therefore, SCP requires efficient recovery of the extremely limited amounts of proteins and peptides available from individual cells while minimizing sample loss, together with highly sensitive detection. In particular, for low-input samples, losses caused by nonspecific adsorption to container surfaces and sample-transfer steps become proportionally more significant, making optimization of each step from sample preparation to LC–MS analysis essential (3, 4). To minimize sample loss, various sample-preparation methods have been developed in which reaction volumes are reduced to the nanoliter or sub-microliter scale. Approaches such as nanoPOTS, nPOP, and proteoCHIP enable single-cell isolation and sample preparation in miniaturized reaction volumes through the use of microfabricated devices or high-precision low-volume dispensing systems such as cellenONE (3, 5, 6). Although these approaches are effective in reducing sample loss, they may require specialized microfabricated devices or high-precision dispensing systems, potentially increasing the cost and complexity of implementation and routine operation. More recently, SCP workflows that reduce dependence on specialized low-volume dispensing devices, for example by combining fluorescence-activated cell sorting (FACS) with conventional multiwell plates, have also been developed (7). In this context, we previously developed single-cell proteomics by LMNG-assisted sample preparation (scpLASP), which employs the nonionic surfactant lauryl maltose neopentyl glycol (LMNG) and enables low-input and single-cell sample preparation at the microliter scale using conventional laboratory vessels without requiring specialized low-volume dispensing devices (8).

Further improvements in SCP performance require optimization not only of cell lysis and protein digestion but also of the entire downstream analytical workflow, including peptide reconstitution after digestion, LC separation, electrospray ionization, and MS acquisition. In particular, for picogram-level low-input samples, peptide losses due to nonspecific adsorption to container surfaces after digestion can represent a substantial fraction of the available peptide material, and reconstitution conditions can therefore have a major impact on analytical sensitivity (8). Addition of surfactants to the reconstitution solution can reduce such peptide losses; however, the peptide recovery performance of combinations of multiple surfactants has not been systematically evaluated. The composition of the LC mobile phase is another important factor affecting detection sensitivity for low-input samples. Acidic additives used in the mobile phase influence electrospray ionization efficiency, while dimethyl sulfoxide (DMSO) is known to affect peptide ionization efficiency and precursor charge-state distributions (9–11). However, the effects of these mobile-phase conditions on ionization efficiency and identification performance in picogram-level samples remain insufficiently characterized.

More recently, the introduction of the Orbitrap Astral series has substantially improved both proteome depth and acquisition speed in label-free SCP, enabling the identification of thousands of proteins from single-cell samples at high analytical throughput (12–14). As MS acquisition has become faster, the number of cells that can be analyzed per day has emerged as an important performance metric in addition to analytical sensitivity.

However, increasing throughput by shortening the LC gradient may affect chromatographic separation, peak sampling, identification depth, and quantitative reproducibility. Therefore, practical high-throughput SCP requires reducing analysis time while maintaining proteome depth and quantitative reproducibility.

In this study, we aimed to establish a sensitive and high-throughput workflow for low-input and single-cell proteomics by building on our previously developed scpLASP platform and optimizing peptide reconstitution, LC–MS sensitivity, and analytical throughput as an integrated workflow. We systematically evaluated surfactant compositions for peptide reconstitution and LC mobile-phase conditions and assessed their effects on analytical performance for low-input samples. We further shortened the LC gradient from 11 to 5.5 min and evaluated the resulting effects on identification depth and quantitative reproducibility. Finally, we integrated these optimized conditions into a complete workflow and applied it to HEK293-derived Expi293F single-cell proteomics to evaluate its analytical performance.

## 2. Experimental Procedures

### 2.1 Preparation of HEK293T and HeLa Peptide Samples

HEK293T cells were cultured and harvested as previously described (15). Protein extraction and digestion were performed using the SP3-LASP method as previously described (16). The resulting peptides were desalted using SDB-GL-Tips (Cat. No. 7820-11200, GL Sciences Inc., Tokyo, Japan), and peptide concentrations in the eluates were determined using a Fluorescent Peptide Assay Kit (Thermo Fisher Scientific, Waltham, MA, USA). Aliquots containing 100 or 10 ng of peptides were prepared and dried by centrifugal evaporation. The dried peptides were reconstituted in 100 µL of 0.1% trifluoroacetic acid (TFA) containing either 2% acetonitrile (ACN), 0.02% of an individual surfactant, or a combination of two surfactants at 0.01% each (Table 1), yielding nominal peptide concentrations of 1 ng/µL or 100 pg/µL, respectively. For HEK293T analyses, 1 µL of each sample was injected, corresponding to peptide inputs of 1 ng and 100 pg, respectively.

**Table 1.** Surfactants used for the optimization of peptide reconstitution conditions.

| Surfactant name | Abbreviation | Molecular weight | CAS No. |
| --- | --- | --- | --- |
| n-Dodecyl-β-D-maltoside | DDM | 510.62 | 69227-93-6 |
| Trehalose C12 | TC12 | 524.6 | 64622-91-9 |
| 3-Oxatridecyl-α-D-mannoside | 3ODM | 364.47 | 914802-92-9 |
| Decyl Maltose Neopentyl Glycol | DMNG | 949.08 | 1257852-99-5 |
| Lauryl Maltose Neopentyl Glycol | LMNG | 1005.19 | 1257852-96-2 |
| Octyl Glucose Neopentyl Glycol | OGNG | 568.69 | 1257853-32-9 |

A commercially available HeLa peptide digest (20 µg; Cat. No. 88329, Thermo Fisher Scientific) was prepared in 0.1% TFA containing 0.02% DDM at a peptide concentration of 5 ng/µL for the HeLa-based LC–MS optimization experiments. For HeLa analyses, injection volumes of 0.02 and 0.05 µL were used, corresponding to peptide inputs of 100 and 250 pg, respectively.

### 2.2 Sample Preparation for SCP

Sample preparation for SCP was performed using a modified version of our previously reported scpLASP method (8). To prepare collection plates for fluorescence-activated cell sorting (FACS), 8 μL of a solution containing 0.2% trifluoroacetic acid (TFA) and 0.1% lauryl maltose neopentyl glycol (LMNG) was added to each well of a 96-well PCR plate (Cat. No. 0030128664, Eppendorf, Hamburg, Germany). The plates were sealed with plate seals (Cat. No. 5010-21951, GL Sciences Inc., Tokyo, Japan) and stored at −80 °C until use.

Expi293F cells (Thermo Fisher Scientific) were cultured in protein-free FreeStyle 293 Expression Medium in suspension. Cells were harvested by centrifugation and washed with fresh culture medium. Viable cells were sorted using an FP7000 cell sorter (Sony Corporation, Yokohama, Japan). Single cells or 20 cells were sorted into collection plates that had been brought to room temperature. Blank controls (0 cells) were prepared without cell deposition and processed in parallel with the single-cell and 20-cell samples using the same sample-preparation procedure. The plates were then sealed and stored at −80 °C until use.

The plates containing blank, single-cell, or 20-cell samples were brought to room temperature and centrifuged at 2,000 × g for 1 min. Subsequently, 2 μL of 500 mM Tris-HCl (pH 8.0) containing 20 mM CaCl₂ and 1 μL of 10 ng/μL Trypsin Platinum (Cat. No. VA9000; Promega Corporation, Madison, WI, USA) prepared in 0.3% acetic acid were added to each well. The plates were sonicated using a Bioruptor II (Sonic Bio Co., Ltd., Samukawa, Kanagawa, Japan) at the low setting for 5 min, centrifuged at 2,000 × g for 1 min, and incubated at 37 °C for 18 h. Following digestion, the plates were centrifuged at 2,000 × g for 1 min, and 2 μL of 5% TFA was added to each well. The plates were then sonicated again under the same conditions and centrifuged at 2,000 × g for 1 min. Peptides were desalted using SDB-GL-Tips (GL Sciences Inc), eluted with 15 μL of 23% 1-propanol, and dried by centrifugal evaporation. The dried peptides were reconstituted in 5 μL of 0.1% TFA containing 0.01% n-dodecyl-β-D-maltoside (DDM) and 0.01% LMNG, followed by mixing for 15 min. Finally, 4.5 μL of each sample was subjected to LC– MS/MS analysis.

### 2.3 NanoLC-MS/MS

LC–MS/MS analyses were performed using an Orbitrap Astral mass spectrometer (Thermo Fisher Scientific) coupled to a Vanquish Neo UHPLC system (Thermo Fisher Scientific). A FAIMS Pro Duo interface (Thermo Fisher Scientific) was used for measurements with FAIMS, whereas an InSpIon system (AMR, Tokyo, Japan) (17) was used for measurements without FAIMS.

For Orbitrap Astral analyses using a 22-min LC gradient, peptides were separated on a C18 capillary column (25 cm × 75 μm i.d., 1.6 μm particle size; MicrOmics Technologies, Orem, UT, USA). Mobile phase A consisted of 0.1% formic acid in distilled water, and mobile phase B consisted of 0.1% formic acid in 80% acetonitrile. Peptides were separated using the following gradient: 1% B at 800 nL/min for 0–0.4 min; 1–10% B at 800–400 nL/min for 0.4–2.4 min; 10–32% B at 400 nL/min for 2.4–15.4 min; 32–48% B at 400 nL/min for 15.4–19.4 min; 48–98% B at 400 nL/min for 19.4–20.2 min; 98% B at 400–800 nL/min for 20.2–21.7 min; and 98% B at 800 nL/min for 21.7–22.0 min. MS1 spectra were acquired over an *m/z* range of 380–980 at a resolution of 240,000 using the Orbitrap analyzer, with an automatic gain control (AGC) target of 300% and a maximum injection time of 5 ms. MS2 spectra were acquired over an *m/z* range of 150–2,000 using the Astral analyzer, with an AGC target of 500%, a maximum injection time of 3.5 ms, and a normalized collision energy of 25%. The MS2 isolation width was set to 3 Th with window placement optimization. For measurements using FAIMS, the compensation voltage was set to −35 V. Apart from the use of the FAIMS Pro Duo interface or the InSpIon system, all other LC and MS acquisition parameters were identical between the two configurations.

For Orbitrap Astral analyses using an 11-min LC gradient, reconstituted peptides were injected directly onto an Aurora Rapid capillary column (5 cm × 75 μm i.d., 1.7 μm particle size; IonOpticks, Collingwood, VIC, Australia) maintained at 60 °C. For the comparison of mobile-phase acids, mobile phase A consisted of distilled water containing either 0.1% formic acid or 0.5% acetic acid. For DMSO optimization, 0.5% acetic acid was supplemented with 0%, 1%, 3%, or 5% DMSO. Mobile phase B consisted of 80% acetonitrile containing the corresponding additives. Peptides were separated using the following gradient: 1% B at 800 nL/min for 0–0.4 min; 1–10% B at 800–400 nL/min for 0.4–1.4 min; 10–34% B at 400 nL/min for 1.4–7.7 min; 34–48% B at 400 nL/min for 7.7–8.7 min; 48–98% B at 400 nL/min for 8.7–9.2 min; 98% B at 400 nL/min for 9.2–10.2 min; 98% B at 400–800 nL/min for 10.2–10.7 min; and 98% B at 800 nL/min for 10.7–11.0 min. All measurements using the 11-min gradient method were performed with a FAIMS Pro Duo interface at a compensation voltage of −35 V. MS1 spectra were acquired over an *m/z* range of 530–730 at a resolution of 240,000 using the Orbitrap analyzer, with an AGC target of 500% and a maximum injection time of 100 ms. MS2 spectra were acquired over an *m/z* range of 150–2,000 using the Astral analyzer, with an AGC target of 800%, a maximum injection time of 24 ms, and a normalized collision energy of 25%. The MS2 isolation width was set to 8 Th with window placement optimization.

For analyses using an Orbitrap Astral Zoom mass spectrometer (Thermo Fisher Scientific) equipped with a FAIMS Pro Duo interface and coupled to a Vanquish Neo UHPLC system (Thermo Fisher Scientific), reconstituted peptides were injected directly onto an Aurora Rapid XT capillary column (8 cm × 75 μm i.d., 1.7 μm particle size; IonOpticks) maintained at 50 °C and separated using either an 11-min or a 5.5-min LC gradient. Mobile phase A consisted of distilled water containing either 0.1% formic acid, 0.5% acetic acid, or 0.5% acetic acid with 1% DMSO, and mobile phase B consisted of 80% acetonitrile containing the corresponding additive.

The 11-min LC gradient was programmed as follows: 6% B at 800 nL/min at 0 min; 6–10% B at 800 nL/min for 0–0.2 min; 10–15% B at 800–450 nL/min for 0.2–1.2 min; 15% B at 450–400 nL/min for 1.2–1.3 min; 15–28.5% B at 400 nL/min for 1.3–7.8 min; 28.5–45% B at 400 nL/min for 7.8–9.0 min; 45–99% B at 400 nL/min for 9.0–9.4 min; 99% B at 400 nL/min for 9.4–9.8 min; 99% B at 400–800 nL/min for 9.8–10.2 min; and 99% B at 800 nL/min for 10.2–11.0 min.

The 5.5-min LC gradient was programmed as follows: 6% B at 1,200 nL/min at 0 min; 6–10% B at 1,200 nL/min for 0–0.2 min; 10–15% B at 1,200–750 nL/min for 0.2–0.7 min; 15% B at 750–500 nL/min for 0.7–0.8 min; 15–32% B at 500 nL/min for 0.8–4.3 min; 32–45% B at 500 nL/min for 4.3–5.0 min; 45–99% B at 500–1,200 nL/min for 5.0–5.2 min; and 99% B at 1,200 nL/min for 5.2–5.5 min.

For the Orbitrap Astral Zoom methods, the FAIMS compensation voltage was set to −42 V. MS1 spectra were acquired over an *m/z* range of 400–800 at a resolution of 240,000 using the Orbitrap analyzer, with an AGC target of 500% and a maximum injection time of 100 ms. MS2 spectra were acquired over an *m/z* range of 150–2,000 using the Astral analyzer, with an AGC target of 800%, a maximum injection time of 20 ms, and a normalized collision energy of 25%. The MS2 isolation width was set to 10 Th with window placement optimization. For Orbitrap Astral Zoom acquisition, low-input mode was enabled, whereas pre-accumulation was disabled.

A total of 60 single-cell LC–MS/MS runs were attempted. Due to a sample loading issue in which the sample solution was not aspirated during one run, that run was excluded from subsequent analysis; as a result, the final analysis included 59 single-cell runs.

### 2.4 Data analysis

The data-independent acquisition mass spectrometry (DIA-MS) data were analyzed using DIA-NN v2.5.1 Enterprise (https://github.com/vdemichev/DiaNN) (18). An in silico human spectral library was generated from the UniProt human proteome database (proteome ID: UP000005640; 20,659 entries; downloaded on April 4, 2026). The spectral library generation parameters were as follows: protease, Trypsin for SCP analyses and

Trypsin/P for non-SCP analyses; missed cleavages, 2; peptide length, 7–35 amino acids; precursor charge, 2–4; precursor *m/z*, 380–980; and fragment *m/z*, 150–2,000. For *in silico* spectral library generation, “Predicting from FASTA” was enabled. “N-term M excision” was enabled for all analyses, whereas “Carbamidomethyl (C)” was enabled for non-SCP analyses and disabled for SCP analyses. DIA-NN searches were performed using the following parameters: MS1 accuracy, 4 ppm; MS2 accuracy, 8 ppm; scoring, generic; proteotypicity, isoform IDs; machine learning, NNs (cross-validated); and quantification strategy, Legacy (direct). “Protein inference” was enabled for all analyses. “Knowledge Base” was enabled for the DIA-NN with KB analysis and disabled for the DIA-NN without KB analysis. For SCP analysis, two spectral-library strategies were used depending on the analysis. In one strategy, single-cell DIA-MS data were analyzed directly against the in silico human spectral library. In the other strategy, the same 20-cell DIA-MS dataset was analyzed separately with the Knowledge Base enabled or disabled to generate two empirical spectral libraries using the in silico human spectral library. Each resulting library was then used to analyze the same single-cell DIA-MS RAW files under the corresponding Knowledge Base setting. Match-between-runs (MBR) and cross-run normalization were disabled for all analyses. The false discovery rate (FDR) threshold was set to 1% at both the precursor and protein group levels, and the final output was filtered using global q-value thresholds of 0.01 at both levels. Protein groups were identified and counted independently for each run. Protein abundances were calculated by aggregating the quantitative values of unique peptides reported by DIA-NN.

The SCP DIA-MS data were additionally analyzed using Spectronaut version 21 (Biognosys AG, Schlieren, Switzerland). Two analysis strategies were used depending on the analysis. In the first strategy, single-cell DIA-MS data were analyzed using the library-free directDIA+ workflow against the UniProt human proteome database (proteome ID: UP000005640; 20,659 entries; downloaded on April 4, 2026). In the second strategy, a 20-cell spectral library was first generated from DIA-MS data acquired from 20-cell samples using the Pulsar search engine against the same UniProt human proteome database, and the resulting spectral library was subsequently used to analyze the single-cell DIA-MS data.

Database searching was performed using Trypsin specificity, allowing up to two missed cleavages and peptide lengths of 7–35 amino acids. The mass tolerances were set to 4 ppm for MS1 and 8 ppm for MS2. Protein inference was performed using IDPicker. For empirical spectral library generation using Pulsar, FDR thresholds were set to 1% at the peptide-spectrum match (PSM), peptide, and protein group levels. For DIA analysis, precursor q-value cutoffs were set to 0.01 at both the run and experiment levels, whereas protein group q-value cutoffs were set to 0.05 at the run level and 0.01 at the experiment level. Quantification was performed at the MS2 level using peak areas with interference correction. For protein-identification analyses, each SCP run was analyzed independently, consistent with the DIA-NN analysis. Unless otherwise specified, default Spectronaut settings were used.

Chromatographic peak characteristics were evaluated using Skyline version 26.1 (19). MS1 extracted-ion chromatograms were generated using an Orbitrap precursor mass analyzer setting with a resolving power of 240,000 at m/z 200. Three precursor isotope peaks were included, and high-selectivity extraction was disabled. Precursor targets and chromatographic peak boundaries were obtained from DIA-NN results using a precursor q-value threshold of ≤0.01. The DIA-NN-derived peak boundaries were imported into Skyline without manual adjustment. Only monoisotopic precursor peaks with start and end times matching the DIA-NN-derived boundaries within 0.02 min and precursor m/z values within 0.01 Th of the reference values were retained. For each experimental condition, precursors detected across all five replicates were independently selected. Precursors with missing, non-finite, or non-positive values for a given metric in any replicate were excluded from the corresponding analysis. FWHM values were converted from minutes to seconds, peak heights were log2-transformed before averaging, and the number of data points across each peak was used without transformation. Mean values were calculated for each RAW file, and condition-level means and standard deviations were calculated from the five replicate measurements.

### 2.5 Statistical analysis and data visualization

Statistical analyses and data visualization were performed using R version 4.6.0 (R Foundation for Statistical Computing, Vienna, Austria) in RStudio version 2026.7.1.147 (Posit Software, PBC, Boston, MA, USA). Unless otherwise indicated, data were summarized using descriptive statistics, including the mean, median, standard deviation, and interquartile range, as appropriate. Protein groups with non-missing, finite, and positive intensity values were considered quantified.

## 3. Results and Discussion

### 3.1. Optimization of the Reconstitution Solution for Low-Input Proteomics

The scpLASP workflow is illustrated in Fig. 1. In scpLASP, excess LMNG and other contaminants are removed after digestion using a reversed-phase StageTip. In SCP, where large numbers of single-cell samples are analyzed sequentially, repeated introduction of cell-derived hydrophobic components and other contaminants into the nanoLC–MS system may lead to their accumulation on the analytical column and spray tip, potentially resulting in deterioration of chromatographic performance and spray stability. A reversed-phase StageTip cleanup step may therefore help maintain stable LC–MS performance during extended sample sequences. However, peptides eluted from the StageTip must subsequently be dried and reconstituted. For low-input samples, peptide losses caused by nonspecific adsorption to container surfaces during these drying and reconstitution steps can have a substantial impact because of the extremely small amount of sample available. The addition of surfactants such as DDM and DMNG to peptide reconstitution solutions has been reported to reduce such losses (7, 8, 13). Therefore, to determine suitable peptide reconstitution conditions for low-input samples, we systematically evaluated different surfactants and combinations of surfactants in the reconstitution solution.

**Figure 1.**
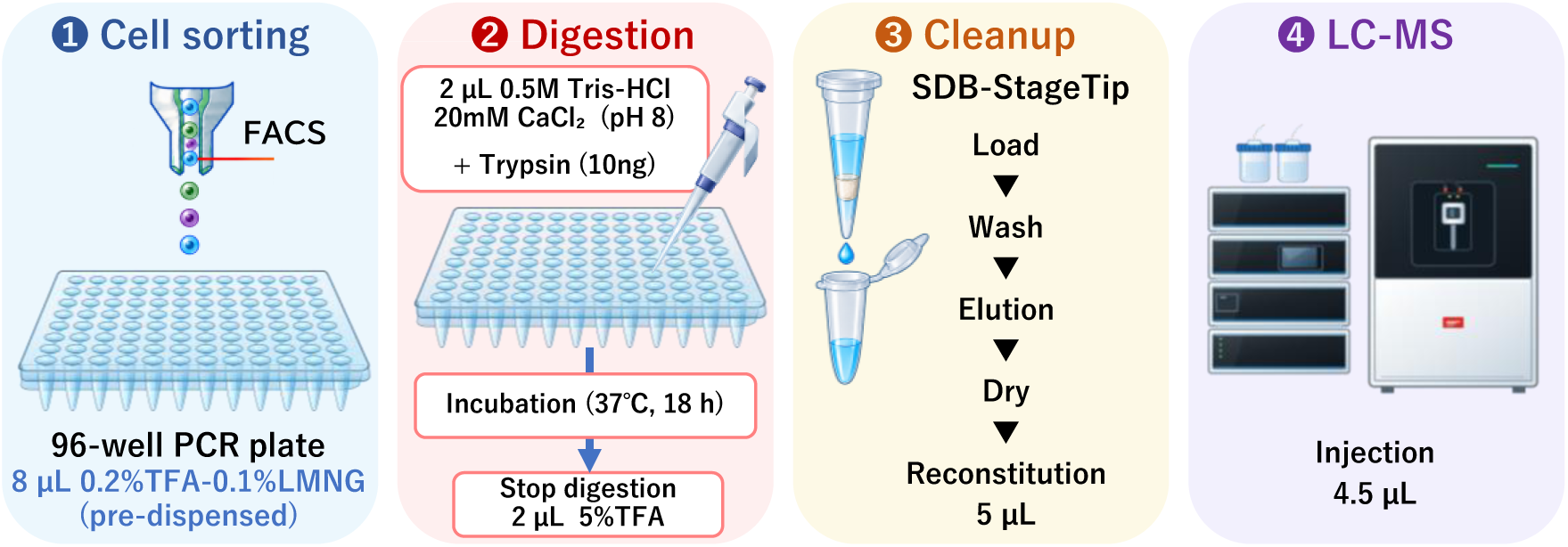
Schematic overview of the scpLASP workflow for single-cell proteomics. Cells were sorted by fluorescence-activated cell sorting (FACS) into individual wells of a 96-well PCR plate preloaded with 8 µL of 0.2% trifluoroacetic acid (TFA) containing 0.1% lauryl maltose neopentyl glycol (LMNG). For protein digestion, Tris-HCl/CaCl₂ buffer and trypsin were added to each well, followed by sonication, centrifugation, and incubation at 37 °C for 18 h. Digestion was terminated by the addition of 5% TFA, followed by a second sonication and centrifugation step. The resulting peptides were desalted using SDB-GL-Tips, dried by centrifugal evaporation, and reconstituted in 5 µL of 0.1% TFA containing 0.01% n-dodecyl-β-D-maltoside (DDM) and 0.01% LMNG. A 4.5-µL aliquot of each reconstituted sample was injected for LC–MS/MS analysis.

First, using 1 ng of HEK293T digest, we compared a total of 22 reconstitution conditions with four replicates per condition (n = 4): six individual surfactants at 0.02%, 15 combinations of two surfactants at 0.01% each, and a surfactant-free control containing 2% ACN and 0.1% TFA (Fig. 2A). The median number of identified peptides was 67,832 under the surfactant-free control condition, whereas higher numbers of identified peptides were observed under many of the surfactant-containing conditions. Among the 22 conditions tested, the combination of 0.01% DDM and 0.01% LMNG yielded the highest median number of identified peptides (82,285 peptides). Compared with the control condition, the number of identified peptides increased by approximately 21.3% with 0.01% DDM and 0.01% LMNG.

**Figure 2.**
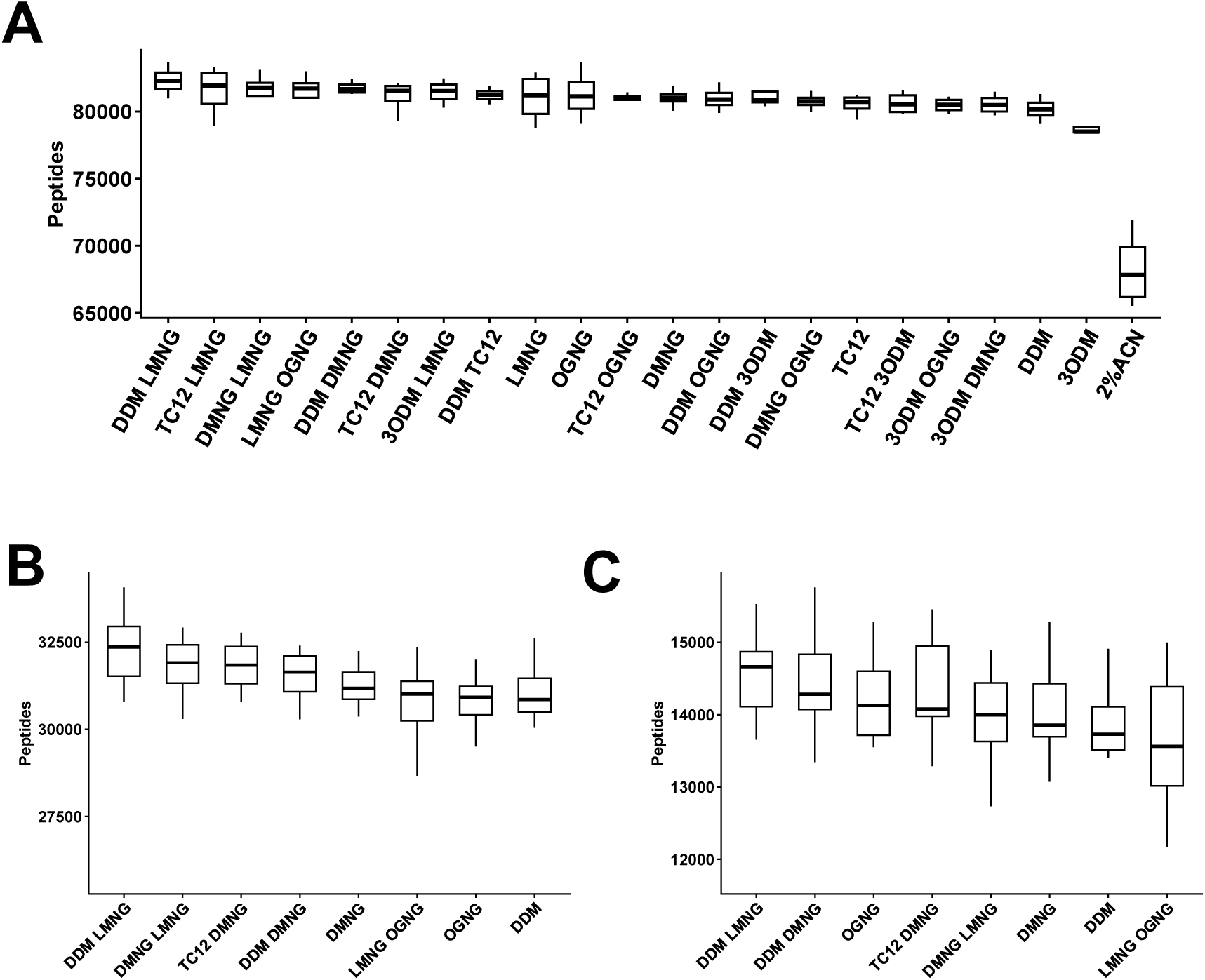
Screening and optimization of surfactant conditions for low-input proteomics. (A) Comparison of peptide identification numbers among individual surfactants, surfactant combinations, and a surfactant-free control using 1 ng of HEK293T digest without FAIMS (n = 4 per condition). (B) Re-evaluation of selected surfactant conditions using 100 pg of HEK293T digest without FAIMS (n = 12 per condition). (C) Evaluation of the same selected surfactant conditions using 100 pg of HEK293T digest with FAIMS (n = 12 per condition). The numbers of identified peptides are presented as box-and-whisker plots. Boxes represent the interquartile range (IQR; 25th–75th percentiles), with horizontal lines indicating the median. Whiskers extend to the most extreme data points within 1.5 × IQR of the lower and upper quartiles. Outliers are not shown. Match-between-runs (MBR) was disabled for all analyses.

Next, the candidate conditions that showed high peptide identification numbers at the 1 ng input level were reevaluated using a reduced sample input of 100 pg (Fig. 2B). At 100 pg, the DDM + LMNG condition again yielded the highest number of identified peptides among the conditions tested, with a median of 32,362 peptides. We further evaluated the same candidate conditions using 100 pg samples analyzed with FAIMS. Under these conditions, DDM + LMNG again showed the highest peptide identification performance, with a median of 14,664 identified peptides (Fig. 2C). These results demonstrate that the superior performance of DDM + LMNG was not limited to the initial 1 ng screening conditions but was also maintained when the sample input was reduced to 100 pg and when FAIMS was used.

Overall, conditions containing combinations of two surfactants tended to yield higher numbers of identified peptides than those containing a single surfactant. Surfactants are known to reduce nonspecific adsorption of peptides to container surfaces, although their effects may vary depending on the physicochemical properties of individual peptides, including hydrophobicity (4). Combining surfactants with different physicochemical properties may therefore reduce adsorption losses for a broader range of peptides, potentially contributing to the higher peptide identification numbers observed with some combinations. Based on these results, 0.01% DDM + 0.01% LMNG in 0.1% TFA was selected for reconstitution of the single-cell samples.

### 3.2. Optimization of Mobile-Phase Additives for Low-Input Proteomics

To improve LC–MS sensitivity and identification performance for low-input samples, we first performed stepwise optimization of mobile-phase additives using an Orbitrap Astral mass spectrometer. Using 100 pg of HEK293T digest, we initially compared 0.1% formic acid (FA), which is commonly used as an acidic mobile-phase additive, with 0.5% acetic acid (AA), which has been reported to improve sensitivity in peptide LC–MS analysis (20, 21). Compared with 0.1% FA, 0.5% AA increased the mean number of identified protein groups from 5,587 to 5,859 (Fig. 3A). The mean numbers of identified peptides and precursors also increased from 28,083 to 31,806 and from 28,109 to 31,843, respectively (Figs. 3B and 3C). In addition, when signal intensities were compared for precursors consistently detected across all replicates under both conditions, 96.2% of the precursors showed increased signal intensity with 0.5% AA (Fig. 3D).

**Figure 3.**
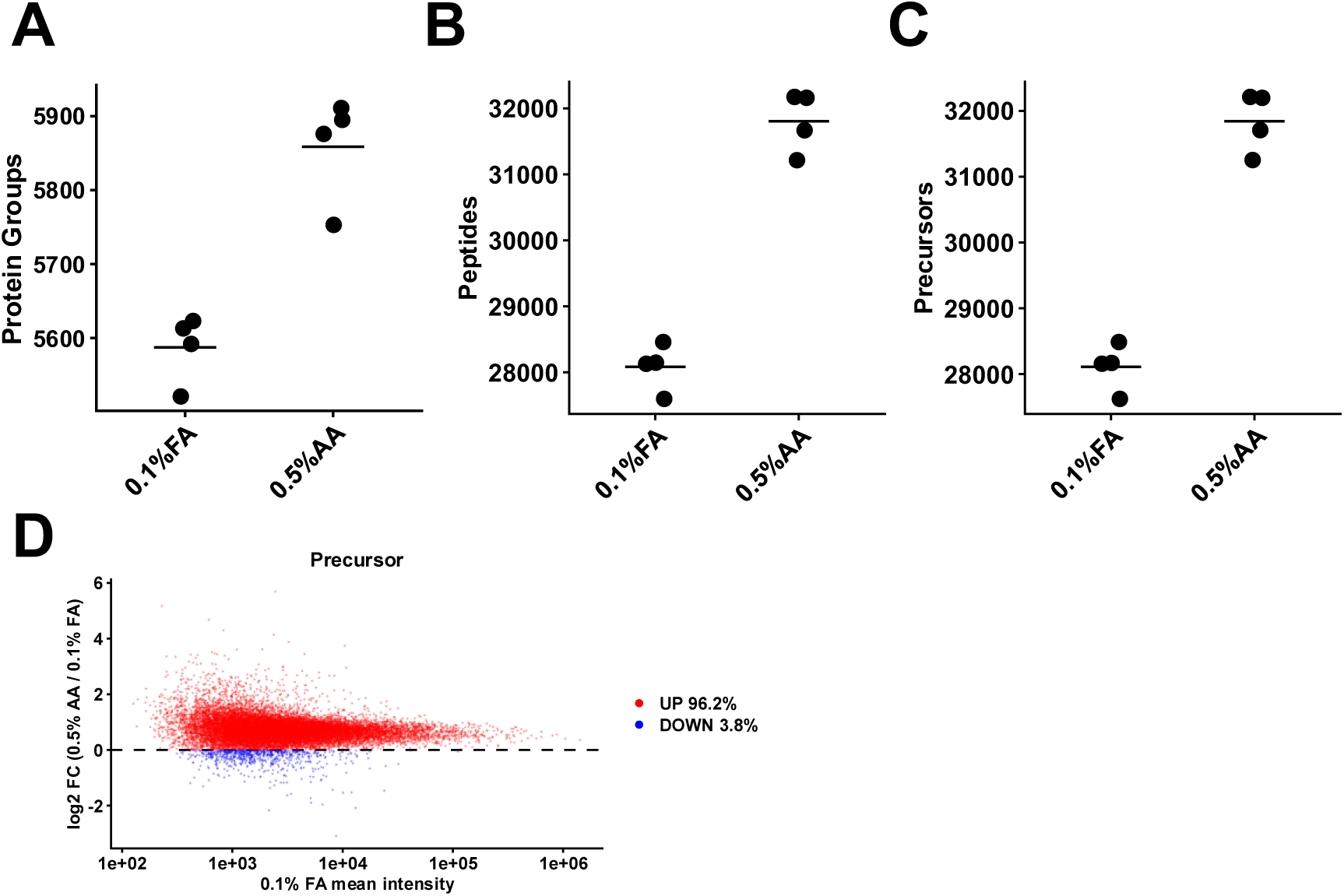
Effect of mobile-phase acid composition on proteomic performance and precursor signal intensity. HEK293T digest (100 pg) was analyzed using an Orbitrap Astral mass spectrometer (n = 4 per condition). (A–C) Numbers of identified (A) protein groups, (B) peptides, and (C) precursors obtained using 0.1% formic acid (FA) or 0.5% acetic acid (AA) as the mobile-phase additive. Each dot represents an individual measurement, and horizontal bars indicate the mean. (D) Comparison of precursor signal intensities between the 0.1% FA and 0.5% AA conditions for precursors quantified across all replicates under both conditions. The x-axis shows the mean precursor intensity under the 0.1% FA condition, and the y-axis shows the log2 fold change in precursor intensity, calculated as the log2-transformed ratio of the mean intensity under 0.5% AA to that under 0.1% FA. Red and blue dots indicate precursors with increased and decreased intensities, respectively, under the 0.5% AA condition. The corresponding percentages are indicated. MBR was disabled for all analyses.

These results are consistent with previous reports demonstrating improved LC–MS sensitivity with AA and indicate that this effect is retained even at a low sample input of 100 pg. Differences in electrospray ionization response between acidic additives may be associated with physicochemical properties of the mobile phase, including ionic strength, surface tension, and ion-pairing characteristics. In particular, reduced mobile-phase ionic strength has been reported to contribute to enhanced signal intensity (22, 23). Based on these results, 0.5% AA was selected as the baseline mobile-phase condition for subsequent optimization.

We next evaluated the effect of DMSO concentration by adding 1%, 3%, or 5% DMSO to the 0.5% AA mobile phase. Although the mean numbers of identified peptides and precursors decreased with increasing DMSO concentration, the mean number of identified protein groups was highest with 0.5% AA + 1% DMSO, increasing from 5,859 with 0.5% AA alone to 6,013 (Figs. 4A–C). Based on these results, 0.5% AA + 1% DMSO was selected as the optimized mobile-phase condition for subsequent experiments.

**Figure 4.**
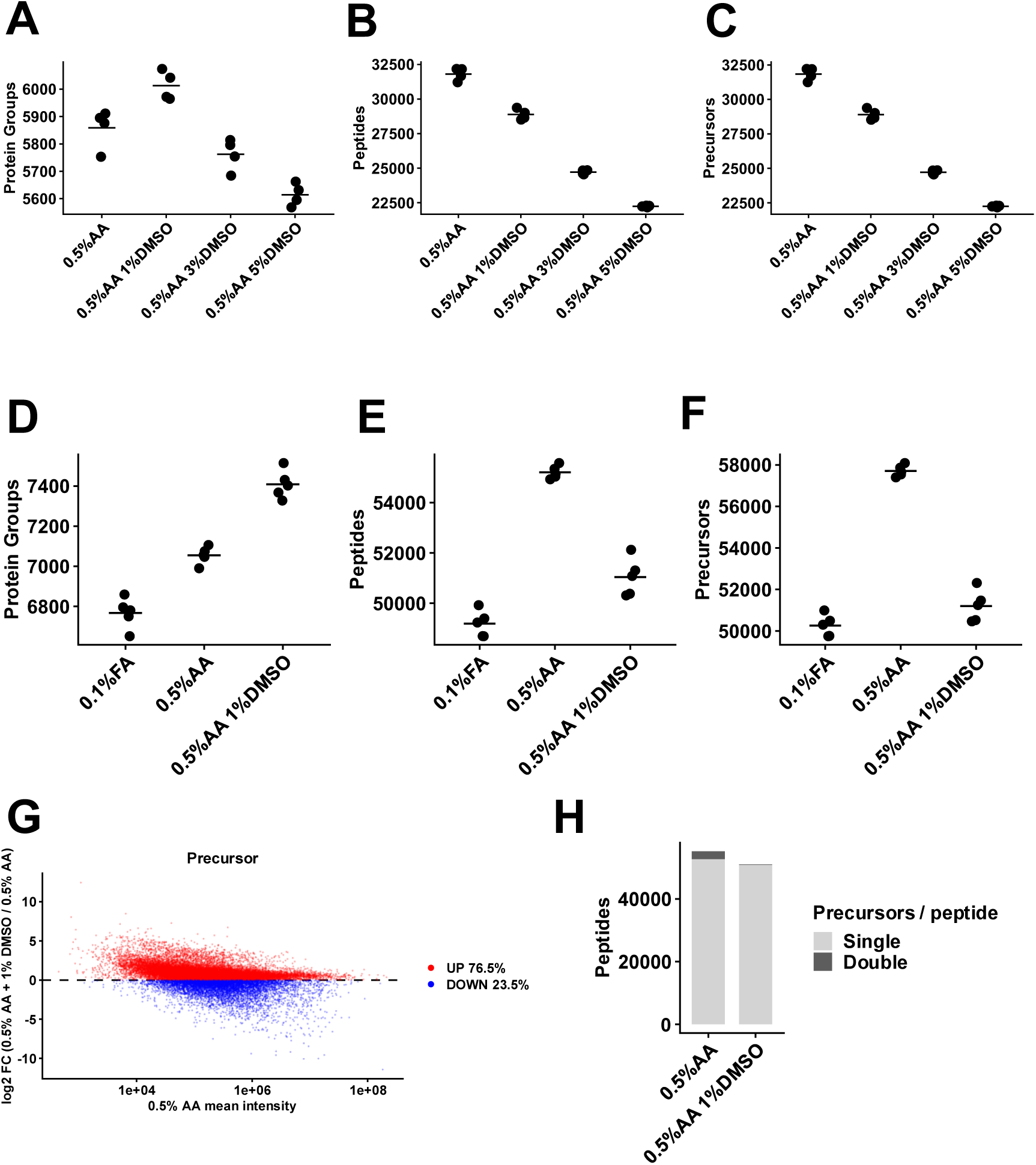
Effect of dimethyl sulfoxide (DMSO) addition to the mobile phase on proteomic performance and precursor characteristics. (A–C) HEK293T digest (100 pg) was analyzed using an Orbitrap Astral mass spectrometer (n = 4 per condition). Numbers of identified (A) protein groups, (B) peptides, and (C) precursors were compared among mobile phases containing 0.5% acetic acid (AA) with 0%, 1%, 3%, or 5% DMSO. (D–H) HeLa digest (250 pg) was analyzed using an Orbitrap Astral Zoom mass spectrometer (n = 5 per condition). (D–F) Numbers of identified (D) protein groups, (E) peptides, and (F) precursors were compared among 0.1% formic acid (FA), 0.5% AA, and 0.5% AA with 1% DMSO. For (A–F), each dot represents an individual measurement, and horizontal bars indicate the mean. (G) Comparison of precursor signal intensities between the 0.5% AA and 0.5% AA with 1% DMSO conditions. The x-axis shows the mean precursor intensity under the 0.5% AA condition, and the y-axis shows the log2 fold change in precursor intensity, calculated as the log2-transformed ratio of the mean intensity under 0.5% AA with 1% DMSO to that under 0.5% AA alone. Red and blue dots indicate precursors with increased and decreased intensities, respectively, following DMSO addition. (H) Distribution of peptides according to the number of detected precursors per peptide under the 0.5% AA and 0.5% AA with 1% DMSO conditions. Peptides represented by one or two detected precursors are indicated separately. MBR was disabled for all analyses.

We next evaluated the effectiveness of the mobile-phase condition selected using the Orbitrap Astral on the higher-sensitivity Orbitrap Astral Zoom platform. For these measurements, a commercially available HeLa digest was prepared in 0.1% TFA containing 0.02% DDM. Using 250 pg of HeLa digest, we compared three mobile-phase conditions: the commonly used 0.1% FA, 0.5% AA, and 0.5% AA + 1% DMSO. The mean number of identified protein groups increased progressively from 6,767 with 0.1% FA to 7,055 with 0.5% AA and 7,409 with 0.5% AA + 1% DMSO (Fig. 4D). These results indicate that the sensitivity improvement associated with AA reported previously was reproduced under low-input conditions using 250 pg of HeLa digest on the Orbitrap Astral Zoom, and that the addition of 1% DMSO further increased protein identification depth. When the three mobile-phase conditions were compared, 0.5% AA yielded the highest numbers of identified peptides and precursors, consistent with the results obtained using the Orbitrap Astral (Figs. 4E and 4F). We next compared signal intensities for precursors consistently detected under both the 0.5% AA and 0.5% AA + 1% DMSO conditions. The addition of 1% DMSO increased the signal intensity of 76.5% of the common precursors, with the increase being particularly pronounced for low-intensity precursors (Fig. 4G). Furthermore, 1% DMSO increased the proportion of peptides represented by a single detected precursor (Fig. 4H).

DMSO is known to enhance ionization efficiency during electrospray ionization and to concentrate peptide ions into a smaller number of charge states (10). In the present study, although the total number of identified precursors decreased following DMSO addition, the number of identified protein groups increased, accompanied by increased signal intensities of common precursors and a shift toward fewer detected precursors per peptide. These observations suggest that the increase in protein group identifications induced by DMSO was not driven simply by an increase in the total number of precursor identifications. Rather, enhanced ionization efficiency and redistribution of peptide ions into fewer charge states may have increased precursor signal intensity and thereby improved the detectability of peptides derived from low-abundance proteins. Notably, these effects were clearly observed even at low sample inputs of 100–250 pg, supporting 0.5% AA + 1% DMSO as an effective mobile-phase condition for increasing protein identification depth in low-input proteomics.

### 3.3. Optimization of LC–MS Throughput and Quantitative Reproducibility

To increase analytical throughput for low-input proteomics, we compared the performance of 11-min and 5.5-min LC gradients using an Orbitrap Astral Zoom mass spectrometer. For 100 pg HeLa digest, the mean numbers of identified protein groups were 7,027 with the 11-min gradient and 6,558 with the 5.5-min gradient, whereas for 250 pg HeLa digest, the corresponding values were 7,417 and 7,088, respectively (Fig. 5A). Thus, although shortening the LC gradient resulted in a modest decrease in the number of identified protein groups, the 5.5-min gradient still identified more than 6,500 protein groups from 100 pg and more than 7,000 protein groups from 250 pg, demonstrating that high proteome depth was largely maintained with the shorter gradient. To further benchmark the analytical performance of our workflow, we reanalyzed a publicly available dataset acquired using an Orbitrap Astral Zoom under the same data-analysis conditions used in the present study (PXD070599) (12). In this dataset, acquired using a 24-min LC gradient (n = 3), the mean numbers of identified protein groups were 6,199 and 6,711 for 100 and 250 pg of HeLa digests, respectively. Although direct comparisons between independent studies should be interpreted with caution because of differences in sample preparation, LC–MS conditions, and other experimental factors, the 5.5-min gradient developed in this study yielded high protein group identification numbers even compared with the previously reported dataset acquired using a 24-min LC gradient. These results demonstrate that the optimized LC–MS workflow can support higher analytical throughput while maintaining high identification depth for low-input samples in the 100–250 pg range.

**Figure 5.**
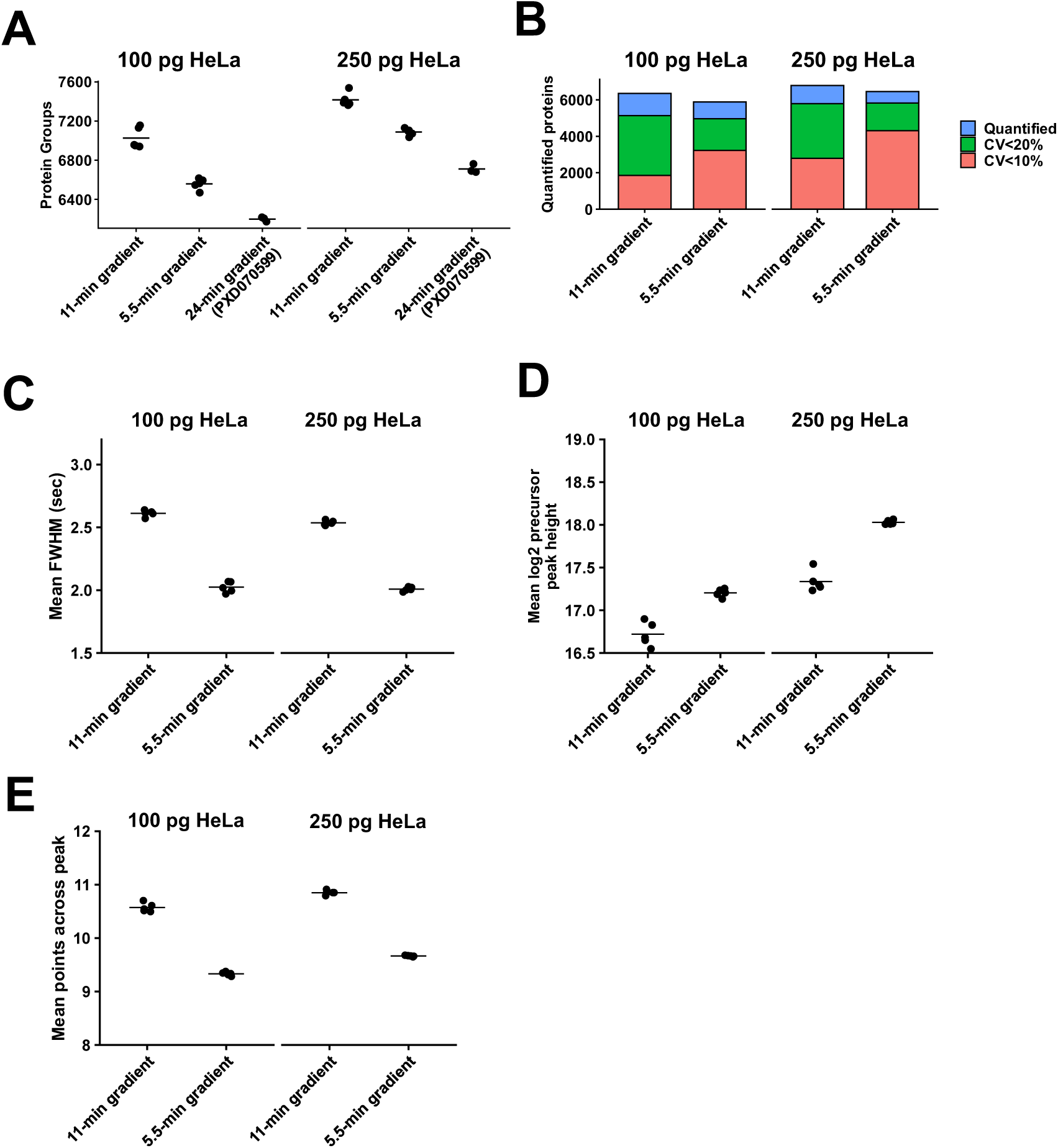
Evaluation of high-throughput low-input proteomics using 11-min and 5.5-min LC gradients. HeLa digests were analyzed using an Orbitrap Astral Zoom mass spectrometer (n = 5 per condition for the 11-min and 5.5-min gradients). (A) Numbers of identified protein groups from 100 pg and 250 pg HeLa digests using the 11-min and 5.5-min gradients. Publicly available Orbitrap Astral Zoom data acquired using a 24-min LC gradient (PXD070599; n = 3) were reanalyzed under the same data-analysis conditions and included for comparison. (B) Protein-level coefficients of variation (CVs) for protein groups quantified across all replicate measurements under the 11-min and 5.5-min gradients. The total height of each stacked bar represents the number of quantified protein groups. The red segment represents protein groups with CV <10%, while the combined red and green segments represent those with CV <20%. (C–E) Chromatographic characteristics of precursor peaks, including (C) mean full width at half maximum (FWHM), (D) mean log2 peak height, and (E) mean number of data points across each peak. Results are shown separately for 100 pg and 250 pg HeLa digests. For (A) and (C–E), each dot represents an individual measurement, and horizontal bars indicate the mean. MBR was disabled for all DIA-NN analyses underlying these results.

We next evaluated whether shortening the LC gradient affected quantitative reproducibility. Protein-level coefficients of variation (CVs) were calculated for protein groups quantified across all replicate measurements within each condition. For 100 pg HeLa digest, the median protein-level CV decreased from 12.34% with the 11-min gradient (n = 6,356 protein groups) to 9.20% with the 5.5-min gradient (n = 5,890 protein groups), while the number of protein groups with CV <10% increased from 1,853 to 3,223 (Fig. 5B). Similarly, for 250 pg HeLa digest, the median CV decreased from 11.01% with the 11-min gradient (n = 6,788 protein groups) to 6.99% with the 5.5-min gradient (n = 6,457 protein groups), and the number of protein groups with CV <10% increased from 2,789 to 4,309. These results indicate that shortening the LC gradient from 11 to 5.5 min did not compromise quantitative reproducibility and that low protein-level CVs were maintained for both 100 pg and 250 pg samples.

To investigate the chromatographic features underlying the favorable quantitative reproducibility observed with the 5.5-min gradient, MS1 chromatograms were analyzed using Skyline (19). For both 100 pg and 250 pg samples, precursors detected in all five replicates within each condition were independently selected and evaluated for full width at half maximum (FWHM), peak height, and the number of points across the chromatographic peak. The mean FWHM decreased from 2.61 s with the 11-min gradient to 2.02 s with the 5.5-min gradient for 100 pg samples and from 2.54 s to 2.01 s for 250 pg samples, indicating that shortening the gradient produced narrower chromatographic peaks (Fig. 5C). Consistent with this peak narrowing, precursor peak heights were approximately 1.4-fold higher with the 5.5-min gradient than with the 11-min gradient for 100 pg samples and 1.6-fold higher for 250 pg samples (Fig. 5D). In contrast, the mean number of points across the peak decreased from 10.6 to 9.33 for 100 pg samples and from 10.8 to 9.67 for 250 pg samples; nevertheless, approximately nine data points per chromatographic peak were maintained even with the 5.5-min gradient (Fig. 5E). These results suggest that the increased peak height associated with narrower chromatographic peaks, together with sufficient data points for peak quantification, may have contributed to the improved quantitative reproducibility observed with the 5.5-min gradient. Thus, shortening the LC gradient not only increased analytical throughput but also maintained favorable chromatographic characteristics for reproducible quantification of low-input samples.

### 3.4. Application of the Optimized Workflow to Single-Cell Proteomics

To evaluate the applicability of the optimized LC–MS workflow to single-cell proteomics, we analyzed Expi293F cells derived from HEK293F cells. Blank controls (0 cells), single-cell samples, and 20-cell samples were prepared using scpLASP. LC–MS analyses were performed using an Orbitrap Astral Zoom mass spectrometer with a 5.5-min LC gradient. With an injection volume of 4.5 µL for single-cell samples, the effective analytical throughput, including sample-loading time, was 135 samples per day (SPD). After exclusion of one single-cell run due to a sample-loading failure, 59 single-cell runs were included in the analysis. Using an *in silico* spectral library, a median of 4,497 protein groups was identified from single-cell samples (n = 59), whereas a median of 8,063 protein groups was identified from 20-cell samples (n = 6) (Fig. 6A). In contrast, no protein groups were identified in four of the six blank controls (0 cells; n = 6), and only 120 and 122 protein groups were identified in the remaining two controls, respectively. When an empirical spectral library generated from 20-cell samples was used, the median number of identified protein groups increased from 4,497 to 5,655 for single-cell samples and from 8,063 to 8,272 for 20-cell samples. Under this condition, no protein groups were identified in two of the six blank controls, while the remaining four controls yielded only 104–185 protein groups. These results demonstrate that the optimized workflow enables deep proteome coverage from individual cells while maintaining low background levels. We further evaluated the effects of data-analysis software and analysis settings on protein identification performance using the same Expi293F single-cell RAW data. Specifically, we compared three analysis configurations: DIA-NN with the Knowledge Base enabled (DIA-NN with KB), DIA-NN with the Knowledge Base disabled (DIA-NN without KB), and Spectronaut. For a consistent comparison, match-between-runs (MBR) was disabled for all analyses, eliminating identification transfer between runs, and precursor and protein group identifications were filtered at a q-value threshold of 0.01 for both DIA-NN and Spectronaut analyses. The Knowledge Base, available in the commercial Enterprise edition of DIA-NN v2.5 and later, is designed to improve identification sensitivity by incorporating information derived primarily from human datasets into PSM scoring. When individual single-cell RAW files were analyzed using an *in silico* spectral library (or the library-free directDIA+ workflow for Spectronaut), the median number of identified protein groups per cell was 4,497 with DIA-NN with KB, 4,111 with DIA-NN without KB, and 3,988 with Spectronaut. In contrast, when empirical spectral libraries generated from 20-cell samples were used, the corresponding median numbers increased to 5,655, 5,554, and 4,710 protein groups, respectively (Fig. 6B). DIA-NN with KB consistently yielded the highest number of identified protein groups under both spectral-library conditions, suggesting that the use of the Knowledge Base contributes to improved protein identification depth in single-cell proteomics. To further assess the overlap and coverage of consistently quantified protein groups across different data-analysis platforms, we compared protein groups quantified in at least 70% of all single-cell samples using empirical 20-cell spectral libraries. DIA-NN with KB quantified 4,917 protein groups, whereas Spectronaut quantified 3,907 protein groups, of which 3,867 were shared between the two software platforms (Fig. 6C). These shared protein groups accounted for 99.0% of those quantified by Spectronaut and 78.6% of those quantified by DIA-NN with KB, indicating that nearly all protein groups consistently quantified by Spectronaut were also consistently quantified by DIA-NN with KB. Furthermore, DIA-NN with KB quantified an additional 1,050 protein groups that were not consistently quantified by Spectronaut, demonstrating broader coverage of protein groups quantified across the single-cell population. Taken together, these results indicate that DIA-NN with KB achieved the highest protein group identification depth among the analysis configurations evaluated and provided broader coverage of consistently quantified protein groups than Spectronaut.

**Figure 6.**
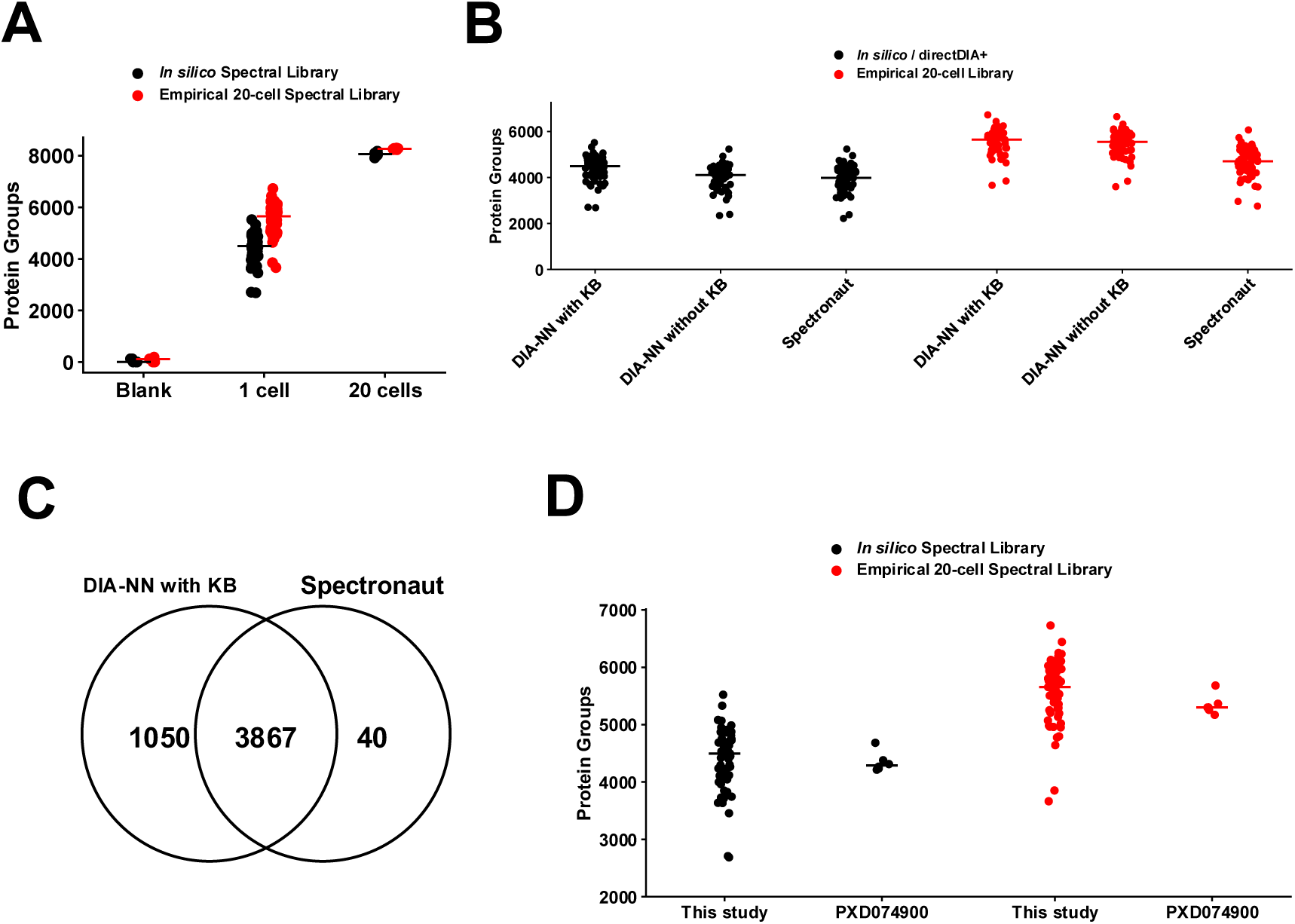
Performance evaluation of the optimized workflow for single-cell proteomics. Expi293F samples containing 0, 1, or 20 cells were prepared using scpLASP and analyzed using an Orbitrap Astral Zoom mass spectrometer. (A) Numbers of identified protein groups in blank (0-cell, n = 6), single-cell (n = 59), and 20-cell (n = 6) samples analyzed using either an *in silico* spectral library or an empirical spectral library generated from 20-cell samples. One of the 60 single-cell runs was excluded due to a sample-loading failure. Each dot represents an individual measurement, and horizontal bars indicate the median. (B) Comparison of protein group identification depth obtained from the same Expi293F single-cell RAW data using DIA-NN with the Knowledge Base enabled (DIA-NN with KB), DIA-NN with the Knowledge Base disabled (DIA-NN without KB), and Spectronaut. Two analysis strategies were evaluated: (i) in silico spectral library searching for DIA-NN or library-free directDIA+ for Spectronaut, and (ii) empirical 20-cell spectral library searching. Each dot represents an individual cell, and horizontal bars indicate the median. (C) Venn diagram showing the overlap of protein groups quantified in at least 70% of Expi293F single-cell samples by DIA-NN with KB and Spectronaut using empirical spectral libraries generated from 20-cell samples. (D) Comparison of the numbers of identified protein groups per single cell between the present study and a previously published HeLa single-cell proteomics dataset (PXD074900, n = 6). Both datasets were reanalyzed using DIA-NN with KB, using either an in silico spectral library or an empirical spectral library generated from the respective 20-cell samples (n = 6 for the present study and n = 3 for PXD074900). Each dot represents an individual cell, and horizontal bars indicate the median. MBR was disabled for all analyses, and precursor and protein group identifications were filtered at a q-value threshold of 0.01 for both DIA-NN and Spectronaut analyses.

To benchmark the performance of the workflow developed in this study against previously reported methods, we reanalyzed a publicly available HeLa single-cell proteomics dataset (PXD074900) acquired by Hao *et al.* (2026) using an Orbitrap Astral Zoom mass spectrometer with a 14-min LC gradient (24). The dataset included six single-cell runs (n = 6) and three 20-cell runs (n = 3), with the latter used to generate the empirical spectral library. The dataset was reanalyzed using DIA-NN with KB, which showed the highest protein identification performance in the preceding comparison, under the same data-analysis criteria applied in the present study. Using an *in silico* spectral library, a median of 4,290 protein groups was identified per cell in the Hao *et al.* dataset, compared with 4,497 protein groups in the present study (Fig. 6D). When empirical spectral libraries generated from the respective 20-cell samples were used, the median number of identified protein groups was 5,300 for the Hao *et al.* dataset and 5,655 for the present study. Thus, the median numbers of identified protein groups were higher in the present study under both spectral-library conditions. Furthermore, whereas Hao *et al.* used a 14-min LC gradient, the present study employed a 5.5-min gradient, representing a reduction of approximately 61%. The sample-preparation strategies also differed: Hao *et al.* employed nanoliter-scale reaction volumes with reagents pre-dispensed using an automated liquid-handling system, whereas the scpLASP workflow used in the present study employs microliter-scale reaction volumes with reagents dispensed using conventional micropipettes. Although direct comparisons of protein identification performance should be interpreted with caution because of differences in cell lines, sample-preparation methods, LC conditions, and MS acquisition parameters, the present study achieved comparable or higher protein group identification numbers while combining a readily accessible sample-preparation method with a substantially shorter LC gradient.

Taken together, these results demonstrate that the optimized LC–MS workflow achieves deep single-cell proteome coverage and high analytical throughput while maintaining low background levels. Among the data-analysis software and configurations evaluated, DIA-NN with KB yielded the highest number of identified protein groups and provided broader coverage of consistently quantified protein groups. Furthermore, comparison with a previously published single-cell proteomics dataset demonstrated that the present workflow achieved comparable proteome coverage despite using a sample-preparation method based on conventional laboratory equipment and a substantially shorter LC gradient. These findings highlight the effectiveness of combining an optimized LC–MS workflow with appropriate data-analysis settings to achieve sensitive and high-throughput single-cell proteomics.

## 4. Conclusion

In this study, we established a sensitive and high-throughput analytical workflow for low-input and single-cell proteomics. To improve peptide identification from low-input samples, peptide reconstitution conditions were systematically evaluated, leading to the selection of 0.01% DDM + 0.01% LMNG. We also optimized the LC mobile-phase composition and found that 0.5% acetic acid + 1% DMSO increased precursor signal intensity and the number of identified protein groups. Using the optimized mobile-phase composition, we shortened the LC gradient from 11 to 5.5 min on the Orbitrap Astral Zoom while maintaining high proteome coverage and favorable protein-level CVs for 100–250 pg HeLa digest samples. When applied to Expi293F single cells, the integrated workflow achieved high identification depth with low background at an analytical throughput of 135 SPD. Comparison with a previously published single-cell proteomics dataset demonstrated comparable proteome coverage despite the substantially shorter LC gradient. Furthermore, comparison of data-analysis software and settings showed that DIA-NN with Knowledge Base provided the highest number of protein group identifications, highlighting the importance of appropriate data-analysis settings for maximizing single-cell proteome coverage.

Taken together, this study establishes a practical workflow that combines low-loss sample preparation at the microliter scale with sensitive and high-throughput LC–MS analysis, enabling deep proteome coverage without reliance on specialized nanoliter-scale sample-preparation devices. This workflow provides an accessible analytical platform for high-sensitivity and high-throughput single-cell proteomics.

## Funding and Additional Information

This work received support from the Kazusa DNA Research Institute Foundation.

## Author Contributions

Y.O., S.W., D.H., and Y.Ka. designed the study; Y.O., S.W., Y.E., R.K., H.M., D.N., and D.H. performed the experiments and contributed to sample preparation and investigation; Y.O. and R.K. analyzed the data; Y.Ko. and O.O. supervised the study and contributed to the interpretation of the results. Y.O. and Y.Ka. wrote the manuscript. All authors have read and approved the final version of the manuscript for publication.

## Data Availability

Mass spectrometry proteomic data were deposited in the ProteomeXchange Consortium through the jPOST partner repository (25, 26).

## Acknowledgements

The authors thank Satoru Yokoyama for technical assistance with fluorescence-activated cell sorting.

